# Co-occurring soil bacteria exhibit a robust competitive hierarchy and lack of non-transitive interactions

**DOI:** 10.1101/175737

**Authors:** Logan M. Higgins, Jonathan Friedman, Hao Shen, Jeff Gore

## Abstract

Microbial communities are typically incredibly diverse, and this diversity is thought to play a key role in community function. However, explaining how this diversity can be maintained is a major challenge in ecology. Temporal fluctuations and spatial structure in the environment likely play a key role, but it has also been suggested that the structure of interactions within the community may act as a stabilizing force for species diversity. In particular, if competitive interactions are non-transitive as in the classic rock-paper-scissors game, they can contribute to the maintenance of species diversity; on the other hand, if they are predominantly hierarchical, any observed diversity must be maintained via other mechanisms. Here, we investigate the network of pairwise competitive interactions in a model community consisting of 20 strains of naturally co-occurring soil bacteria. We find that the interaction network is strongly hierarchical and lacks significant non-transitive motifs, a result that is robust across multiple environments. Moreover, in agreement with recently proposed community assembly rules, the full 20-strain competition resulted in extinction of all but three of the most highly competitive strains, indicating that higher order interactions do not play a major role in structuring this community. The lack of non-transitivity and higher order interactions *in vitro* indicates that other factors, such as temporal or spatial heterogeneity, must be at play in enabling these strains to coexist in nature.

---

Despite their small size, microbes play outsized roles at multiple ecosystem scales, from the planetary^1^ to that of the human individual^2^. Like their macroscopic counterparts, microbes typically exist in diverse communities whose functions are intimately related to their structure. Diversity impacts an ecological community’s stability, resilience to perturbations, and its ability to provide ecosystem services^3^. Therefore, a long-standing area of interest in microbial ecology has been understanding the factors that give rise to the diversity observed within microbial communities^4,5^. A better understanding of the structure of microbial communities is desirable for both managing existing microbial communities^6^ and, eventually, engineering them *de novo^7^*.

Many factors can contribute to the generation and maintenance of diversity in ecological communities. Non-transitivity^8^, bistability^9^, weak interactions^10^, facilitation, multiple limiting factors, and spatial or temporal segregation^11^ have all been hypothesized to play a role; however, there is little empirical data regarding the relative importance of each of these factors in actual natural communities. By investigating the network of underlying interactions among the members of a given community, we can better understand each factor’s relative importance in structuring the community^12^. Since interspecific competition is thought to be a dominant factor in determining whether a given species can persist in a community^13,14^, the network of competitive interactions between species may be particularly informative of the structure of the community within which the interaction takes place. Features of competitive interaction networks that could contribute to community diversity can include non-transitive motifs such as the classic rock-paper-scissors triad, network modularity^15^, or overall trends towards weak interactions among species

While non-transitivity in particular is often cited as a potential driver of interspecies coexistence^16,17,18^, the degree to which it occurs in natural communities remains largely unknown. Indeed, Levine and colleagues recently asserted that despite the theoretical potential of non-transitive interactions to stabilize community structure, there is scant evidence that they are widespread in natural systems, and that further empirical studies are warranted^19^. Recent experimental work using a field-parameterized model of competition in annual plants^20^ and naturally co-occurring *Streptomyces* bacteria^9^ suggest that rock-paper-scissors type interactions may be less common in natural communities than we might assume; however, further studies of competitive interaction networks in diverse ecological communities are warranted, particularly among phylogenetically diverse natural assemblages.

Here, we add to this small but growing body of research that suggests that non-transitive interactions may play a less significant role in maintaining species diversity than is commonly assumed. We use a model system composed of heterotrophic bacteria isolated from a single soil grain. By competing in all pairwise combinations in laboratory culture, we find that the overarching feature of the resulting interaction network is a strong competitive hierarchy, a feature that is naturally at odds with a high incidence of non-transitivity. Therefore, in the natural environment of these bacteria, other factors must be at play that account for their ability to co-occur.

## Results

To probe the network of pairwise interactions in a community of diverse microbes, we isolated a collection 20 strains of naturally co-occurring heterotrophic bacteria from a single grain of soil. This strain collection is phylogenetically diverse and spans 16 species across seven genera and five families (Fig. 1a and Methods). Similar to ref^21^, we co-inoculated all pairwise combinations of the 20 strains at varying initial fractions and propagated them through at least five growth-dilution cycles. During each growth cycle, cells were cultured for 24 hours and then diluted by a factor of 100 into fresh media. The final outcome of competition was determined by plating the cultures on solid agar and counting colonies, which are morphologically distinct (Fig. 1c and Supplementary Fig. 1). Plating results were confirmed via next-generation sequencing for a random subset of the pairs (Supplementary Fig. 6).

**Figure 1.**
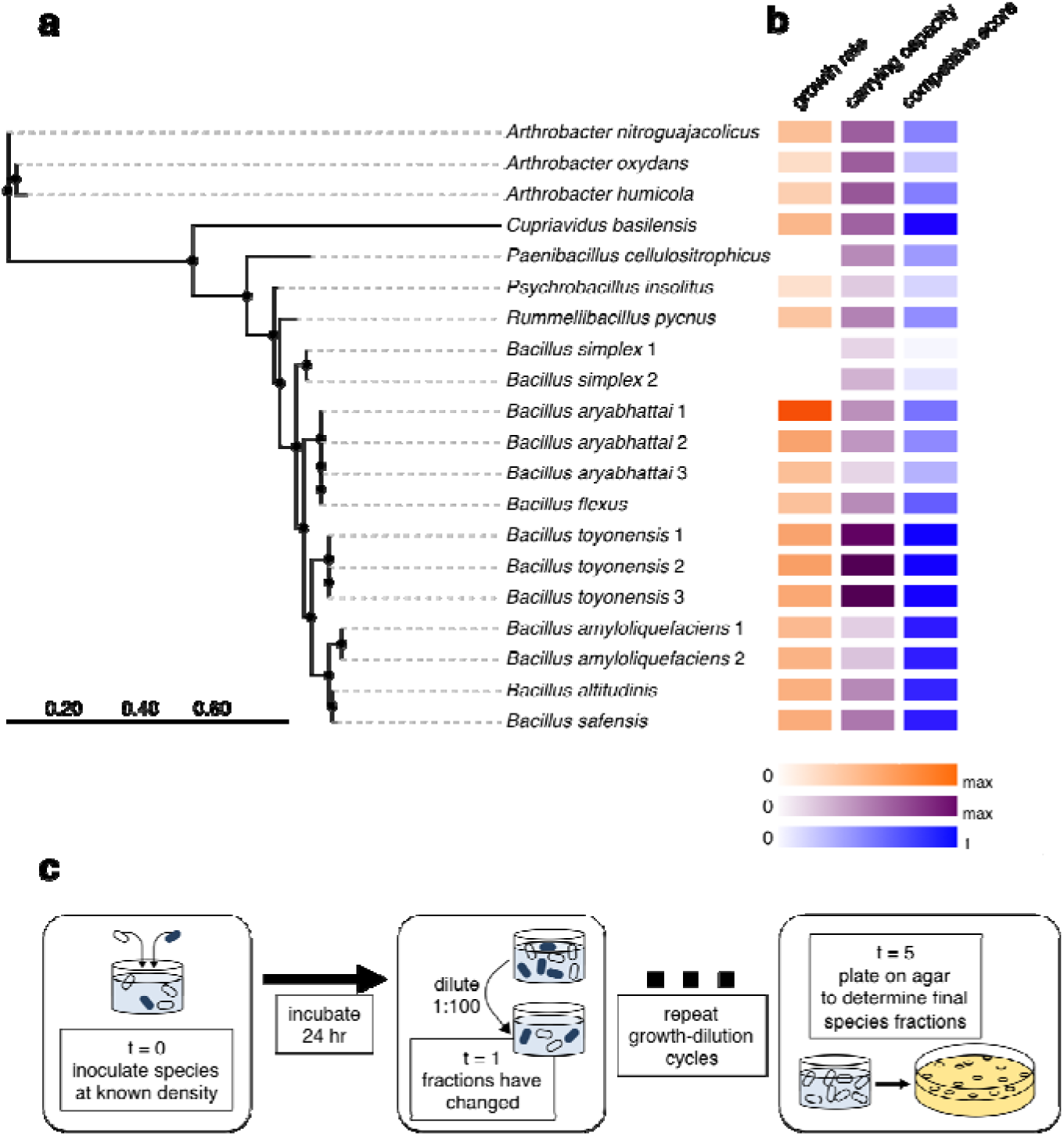
Twenty strains of bacteria isolated from a single grain of soil were competed against each other in all pairwise combinations. **a**, Phylogenetic tree of the 20 strains used in this study. Tree was constructed using the full 16S gene. **b**, Growth rate (orange) and carrying capacity (purple) of each strain in monoculture, as well as competitive score against other strains (blue). Lighter shades correspond to lower values, while darker shades correspond to higher values. **c**, All 190 pairwise combinations of the soil isolates were competed in the laboratory. Colonies of different strains were visually distinct, allowing determination of final species fractions at the end of competition.

Pairwise competitions resulted in one of three qualitatively different outcomes: exclusion, coexistence, or bistability (Fig. 2a-c and Methods). In 153 of the 190 pairs (81%), only one strain could invade the other and drove it to extinction, an outcome we call exclusion. Nineteen pairs (10%) were mutually invasible, and thus exhibited coexistence over the time span of the experiment. Finally, 15 pairs (8%) were mutually non-invasible, an outcome that we call bistability. In a small number of pairs (3; 2%), we were unable to determine the outcome due to contamination. Due to the high incidence of exclusion and bistable outcomes, we conclude that these strains interact in the experimental environment primarily through competition.

**Figure 2.**
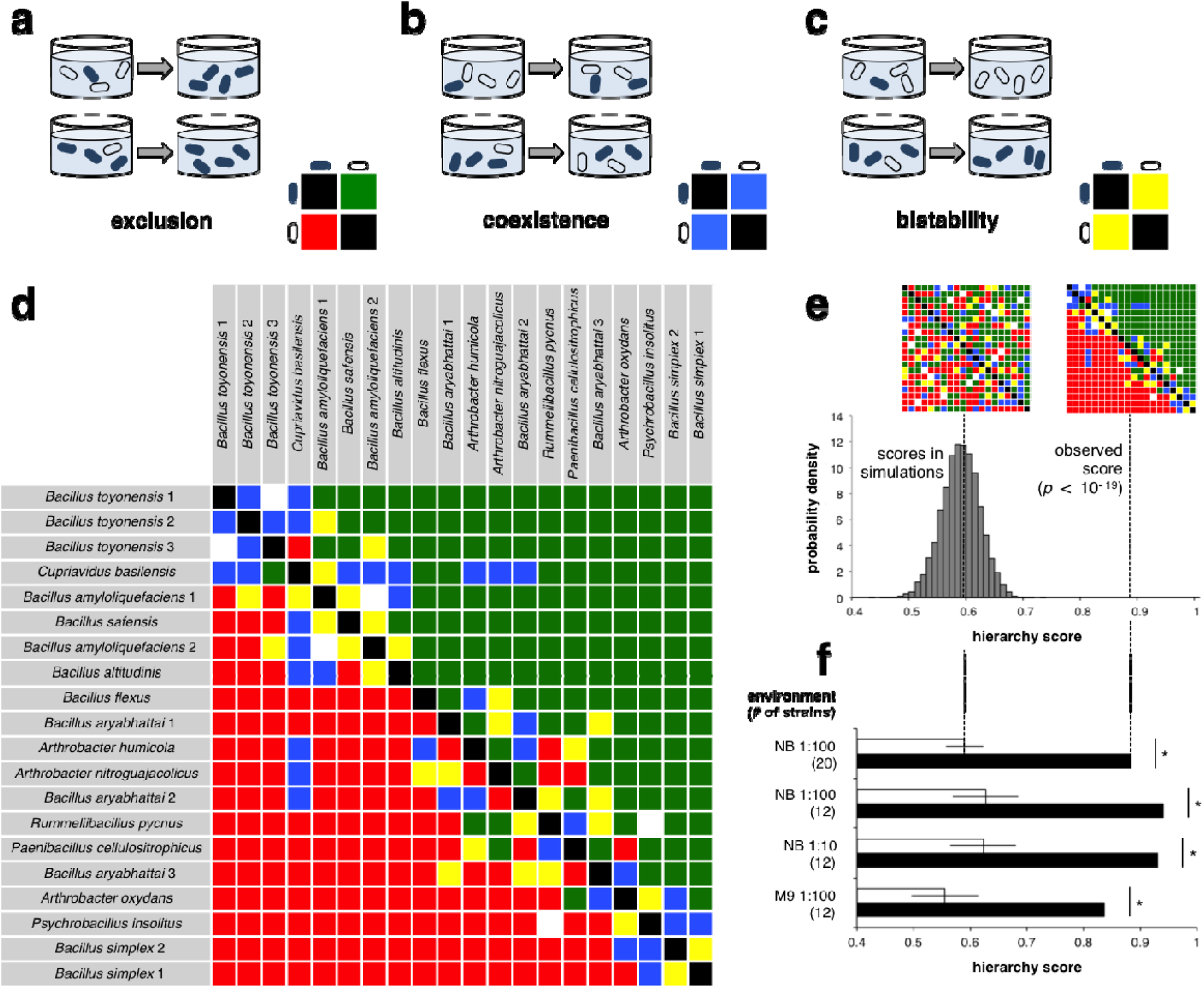
The network of pairwise interactions among strains is strongly hierarchical. **a**-**c**, Changes in relative abundance over time in three hypothetical pairs: one in which the outcome was competitive exclusion; one in which the outcome was stable coexistence; and one in which the outcome was bistability. The color-coded matrices inset into each diagram indicate the qualitative outcome for the row species in competition with the column species. **d**, Pairwise outcome matrix for the entire 20-strain collection. Outcomes are color coded as for **a**-**c**, with white indicating an indeterminate outcome. Rows and columns are sorted in decreasing order of each strain’s competitive score. **e**, Histogram of hierarchy scores for randomized outcome matrices. The hierarchy score for a given matrix is calculated by summing the final fractions of the row strain in competition with the column strain across all row-column pairs in the upper triangle of the matrix. The difference is highly significant (*p* < 10^-20^). **f**, Hierarchy scores for pairwise interaction networks associated with varying environmental conditions and the corresponding randomized networks. NB: 0.2X nutrient broth. M9: 1X M9 minimal medium supplemented with 0.2% casamino acids, 0.4% glycerol, and 1 mM thiamine HCl. Dilution rates were either 1:100 or 1:10 per 24 hr, and experiments consisted of either the full complement of 20 bacterial strains or subsets of 12, as indicated in parentheses. Error bars represent +/– 1 s.d. Differences in observed versus randomized scores were were highly significant in all environments (*p* < 10^-7^).

To quantify the strains’ overall competitive ability, we define each strain’s competitive score to be its mean final fraction across all pairwise competitions. The competitive scores that we measured spanned nearly the entire possible range, from a low of 0.03 to a high of 0.91 (Fig. 1b and Supplementary Table 1).

The strains exhibit a strong competitive hierarchy. Very few strains were able to exclude a strain with a higher competitive score; out of 187 pairwise competitions measured, only five resulted in the lower-ranked strain excluding the higher-ranked one (Fig. 2d). The degree of hierarchy in this interaction network is highly significant when compared to networks with randomized outcomes (*p* < 10^-19^; Fig. 2e). To assess whether the hierarchical pattern was specific to a particular environment, we repeated the competitions with subsets of the full 20-strain collection in different growth media and with different dilution rates (Supplementary Fig. 2). We found that the resultant interaction networks in these different environments were also highly hierarchical, despite changes in which strains were most competitive (Fig. 2f and Supplementary Fig. 3). Thus, we conclude that hierarchy in pairwise competition is a robust feature of this model community.

**Figure 3.**
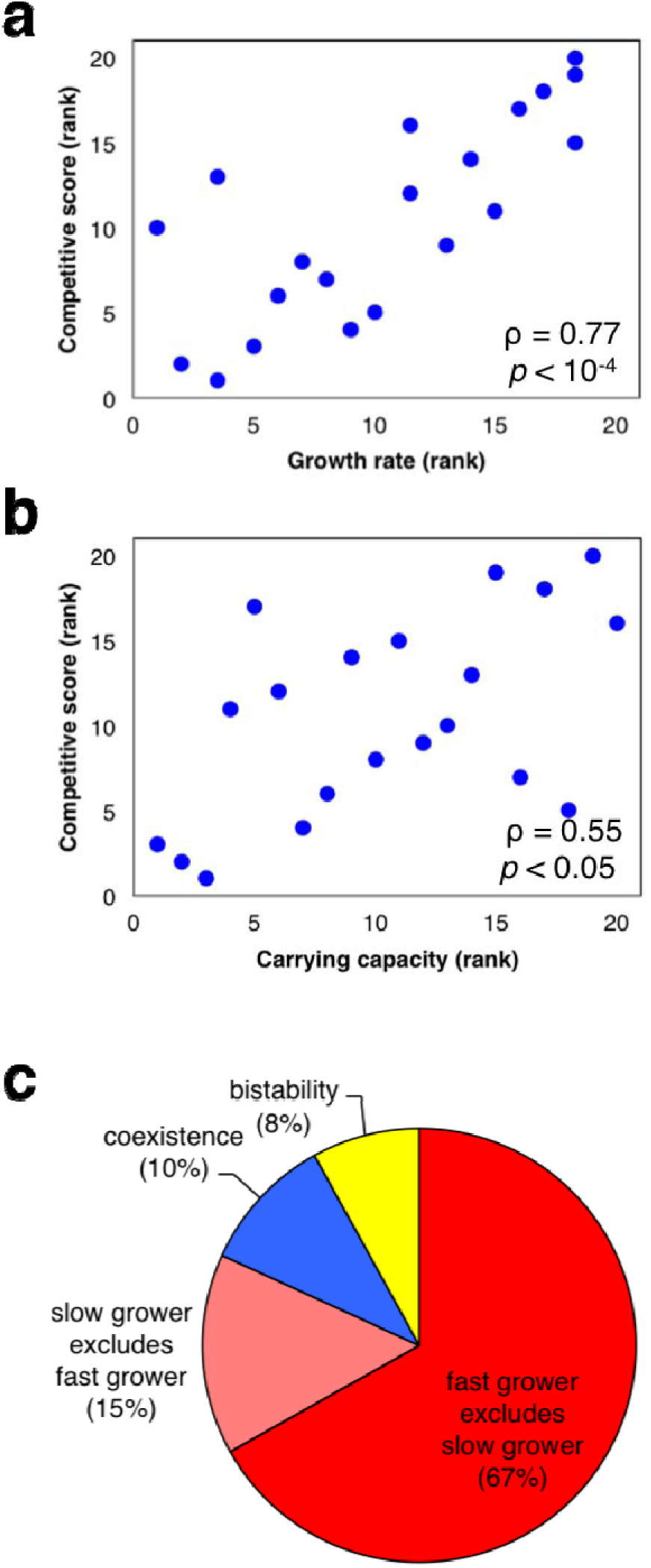
Differences in growth parameters frequently predict the outcome of competition. **a**, **b**, Correlation between rank in growth rate (as estimated using a time-to-threshold method) or rank in carrying capacity (as measured using OD_600_) and rank in competitive score. Figures reported are Spearman correlation coefficients (*ρ*) with two-sided *p*-values. **c**, Distribution of competitive outcomes for all pairs, with pairs that exhibit exclusion differentiated according to whether the faster or slower grower excludes the other.

Next, we asked what characteristics of a strain might best predict its performance in competition. We hypothesized that strains that grow well in monoculture will have competitive advantages over strains that grow more poorly. Indeed, we found that exponential growth rate (*r*) was positively correlated with competitive score (Spearman’s rho = 0.77; *p* < 10^-4^; Fig. 3a) and that the typical outcome was for the strain with the higher *r* to exclude the strain with the lower *r*, which occurred for 67% of pairs (Fig 3c). Carrying capacity (*K*) in monoculture was less predictive of competitive superiority, but was still significantly correlated (Spearman’s rho = 0.55, *p* < 0.05; Fig. 3b). In general, the likelihood of outcomes other than the stronger grower outcompeting the weaker grower decreases for large differences in *r* and *K* (Supplementary Fig. 4). While differences in these two parameters can be indicators of the likelihood of a given competitive outcome, there are many exceptions, and, indeed, some of the stronger competitors do not necessarily have correspondingly strong single-species growth parameters. Thus, while the each species’ intrinsic growth ability correlates with competitive ability, the significant number of exceptions indicates that growth ability alone does not fully explain the hierarchical competitive structure that we observe.

**Figure 4.**
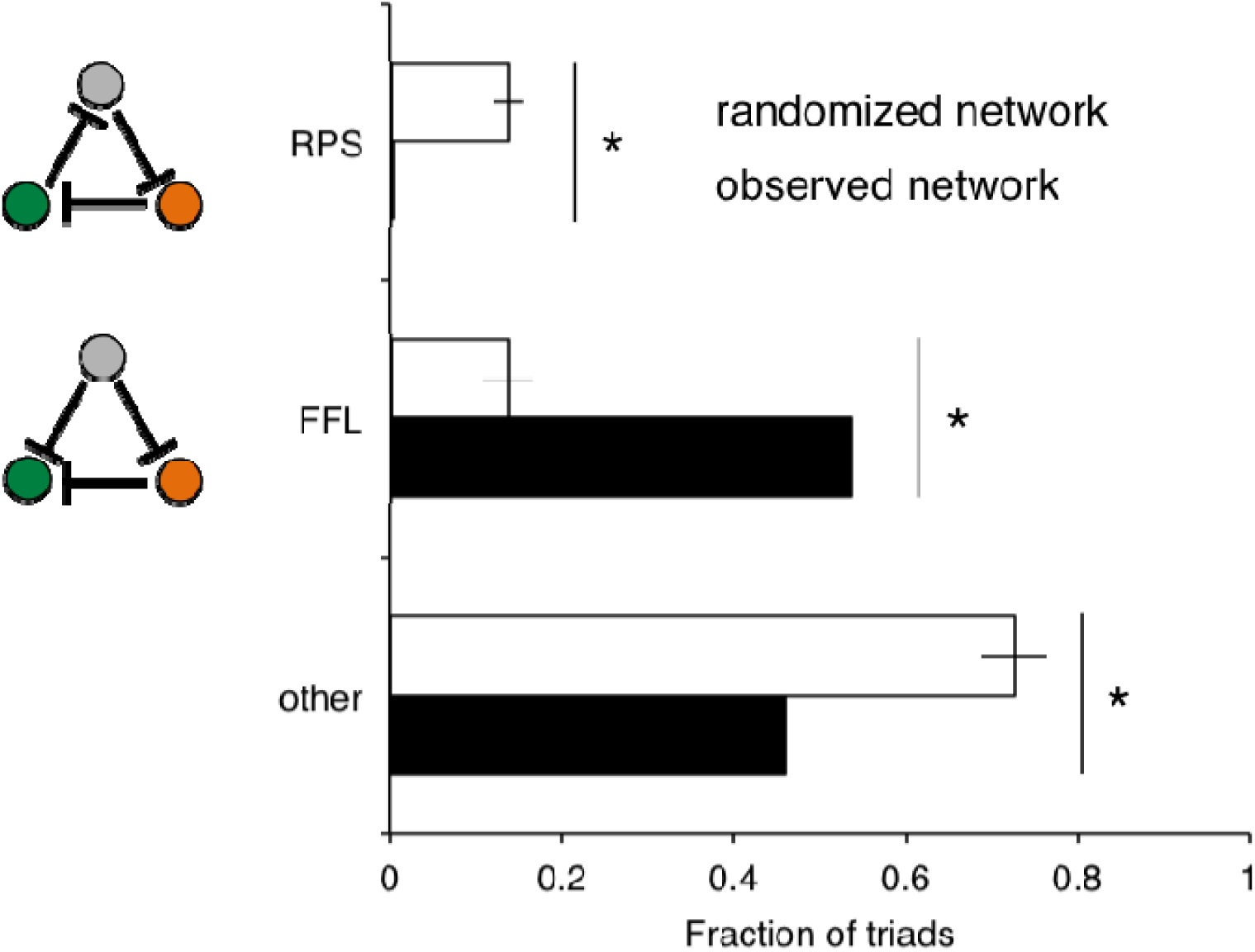
The observed interaction network contains very few cycles. There were significantly fewer rock-paper-scissors triads and significantly more feedforward loops in the network of observed outcomes as compared to 1000 randomized networks. Error bars represent +/– 1 s.d.. Differences in the observed versus randomized incidences were highly significant for all motif categories (*p* < 10^-7^).

An important corollary of the high degree of hierarchy we observed in the interaction network is that non-transitive motifs are vanishingly rare. Non-transitive motifs are instances in which a clear competitive hierarchy among members of a sub-group does not exist, the classic example being a rock-paper-scissors (RPS) triad. Of the 987 triads in our collection for which complete pairwise outcome data are available, only three (0.3%) display the RPS topology. This number is significantly less than is found in randomized networks, where on average 14% of triads were RPS (*p* < 10^15^; Fig. 4). Furthermore, the three triads that we classify as RPS each feature strains that display unusually high variability from experiment to experiment, possibly due to rapid evolution, and further efforts to characterize these triads failed to reproduce the non-transitive network topology. As dictated by its hierarchical structure, our network is also highly enriched for perfectly hierarchical feedforward loops, which were observed in over 50% of triads (Fig. 4). Due to the paucity and irreproducibility of observable non-transitive relationships among our strains *in vitro*, we conclude that such relationships are unlikely to be a significant contributor to their coexistence in a natural environment.

Given the hierarchical structure of the pairwise interaction network, we wondered about the potential of higher-order interactions and indirect effects among our strains to give rise to a diverse community. To address this, we inoculated three replicate cultures with equal proportions of all 20 strains and propagated them through five growth-dilution cycles (Fig. 5b). The resulting assemblages were highly replicable, and consisted of three strains representing some of the strongest competitors in pairwise experiments (Fig. 5a,c), all of which were found to coexist with each other in pairwise competition. Notably, this combination of survivors was consistent with the simple community assembly rule we recently developed^21^: namely, that a strain is expected to survive in multispecies competition if and only if it is not excluded by any other surviving species. Since pairwise outcomes alone are sufficient to predict the outcome of multispecies competition in this environment, we conclude that higher-order interactions are unlikely to play a major role in structuring this community.

**Figure 5.**
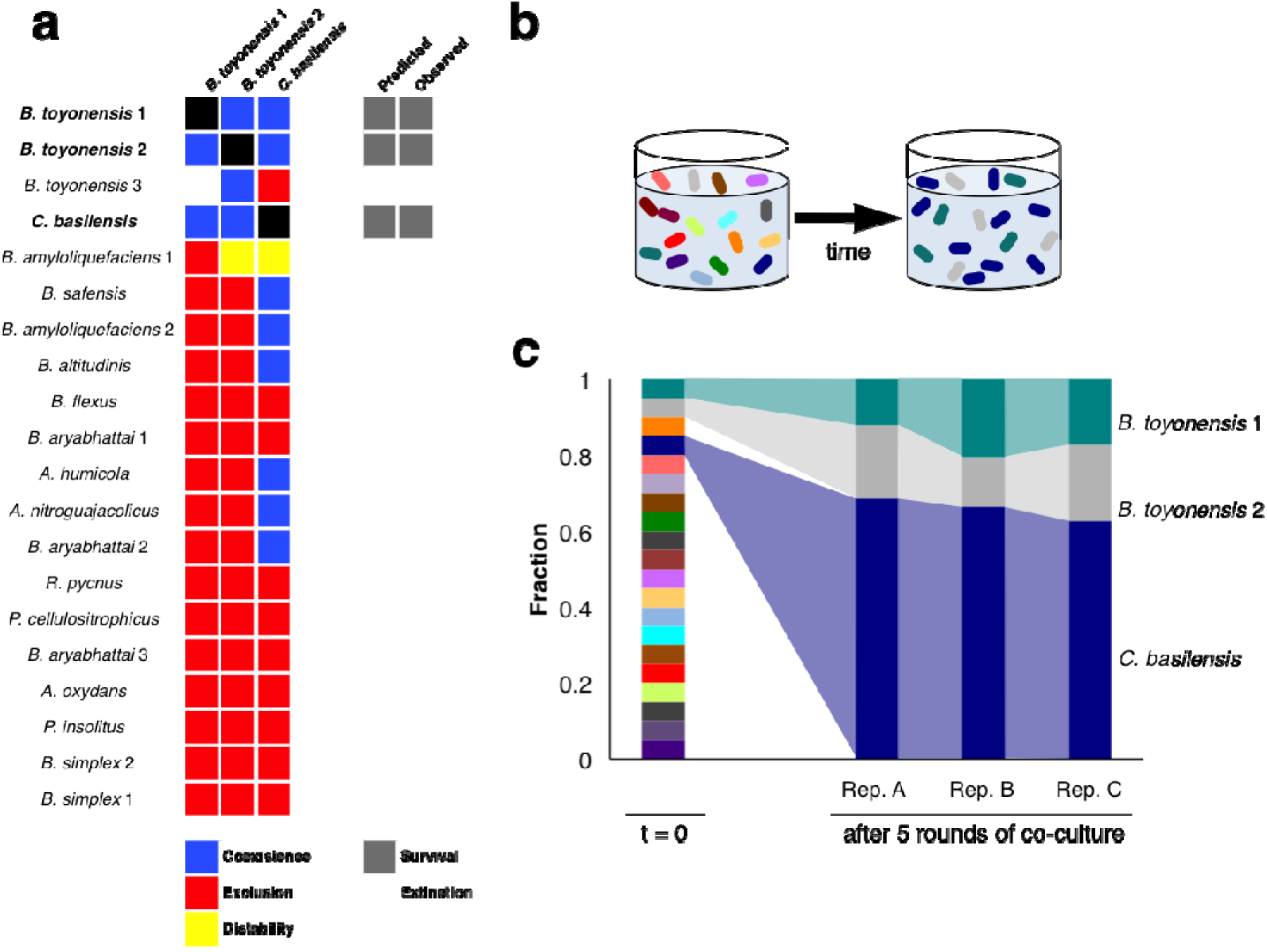
Only three species survive in all-versus-all competition, as predicted by pairwise outcomes. **a**, Predictions and observed outcomes of multispecies competition (grey squares, right) based on community assembly rules incorporating the outcomes of pairwise interactions (colored squares, left). **b**, All strains were mixed in equal proportion by optical density and allowed to reach equilibrium. **c**, In three replicate cultures, only the same three strains survived, each of which was found to coexist with the other two strains in pairwise experiments.

## Discussion

Many factors can contribute to the generation and maintenance of diversity in ecological communities. Non-transitivity, facilitation, bistability, weak interactions, multiple limiting factors, and spatial or temporal segregation have all been hypothesized to play a role^22^; however, there is little empirical data regarding the relative importance of each of these factors in actual natural communities. Here, we explored one such community. In this work, we explored the network of pairwise interactions for a community of naturally co-occurring bacteria. Our results indicate that diversity in this community is likely maintained primarily due to factors including and spatial or temporal segregation or multiple limiting factors, rather than frequent bistability, non-transitivity, or higher order interactions, all of which have been hypothesized to play a role in generating and maintaining diversity. Nonetheless, we still do not completely understand the processes that give rise to the diversity we observe in nature.

Given that soil is a heterogeneous mixture with a multitude of microhabitats, microbial co-occurrence in soil may be facilitated by niche separation and spatial de-mixing. This would allow the coexistence of strains that display strong inhibitory interactions in well-mixed environments. Microbes in soil also experience a strongly fluctuating environment, which can lead to coexistence of multiple strains over time via the soil spore bank. Members of the genus *Bacillus* are particularly well known for their spore-forming ability, which may allow them to persist in a non-vegetative, and therefore non-competitive state, until conditions favor their growth^23^. Finally, our experimental approach clearly requires that the strains to be competed be culturable in the laboratory, so it is impossible for us to exclude the possibility that other strains present within the soil might behave very differently.

Simulations of our experimental system using the generalized Lotka-Volterra model (gLV) predicted that, if the underlying ecological interactions among species are assigned at random, the pairwise interaction network should become less hierarchical at lower death rates, corresponding to a lower daily dilution rate in our experimental setup (Supplementary Fig. 5). In order to test this hypothesis, we competed a subset of pairs while experimentally reducing the dilution rate from 1:100 to 1:10 (Fig. 2f). The hierarchical network structure was robust to this manipulation, and remained highly correlated with growth rates in monoculture. While it is possible that reducing the death rate further could weaken the hierarchy by reducing the importance of a growth rate advantage in determining survival, the most straightforward interpretation of our data is that the hierarchy is not simply due to differences in growth rates.

This experimental system also gives us the opportunity to test the importance of higher order interactions in shaping communities. Higher order interactions are said to take place when the presence of an additional species changes the interaction between two existing species^24^, and have the potential to contribute to the maintenance of species diversity^25^. In bacterial systems, this can be driven by complex networks of selective antibiotic production and sensitivity^26^. Despite the potential for higher order interactions in our model community, our simple assembly rule^21^, which disregards higher order interactions entirely, accurately predicted the survivors in all-versus-all competition *in vitro*, suggesting that higher order interactions are not a major driver of community structure in this instance.

The observation of high levels of diversity in communities of competing organisms is a long-standing paradox in community ecology^27^. In this work, we showed that a bottom-up approach to studying community assembly can be useful in narrowing down the range of possible explanations for the diversity we observe in nature. However, this approach necessitates removing the organisms from their natural environment, including the larger community in which the species of interest are embedded. Future work combining *in vitro* competition experiments with a more mechanistic understanding of the influence of environment on species survival would help to further explain the persistence of diversity in nature.

## Methods

### Strain isolation and identification

Bacterial strains were isolated from a single grain of soil collected in September, 2015 in Cambridge, Mass., U.S.A. The grain weighted ∼1 mg and was handled using sterile technique. The grain was washed in phosphate-buffered saline (PBS) and serial dilutions of the supernatant were plated on nutrient agar (0.3% yeast extract, 0.5% peptone, 1.5% bacto agar) and incubated for 48 hr at room temperature. Isolated colonies were sampled and cultured at room temperature in 5 mL nutrient broth (0.3% yeast extract, 0.5% peptone) for 48 hr. To ensure purity, the liquid cultures of the isolates were diluted in PBS and plated on nutrient agar. Single colonies picked from these plates were once again grown in nutrient broth for 48 hr at room temperature and the resulting stocks were stored in 20% glycerol at -80⁎ C.

The 16S rRNA gene was sequenced via Sanger sequencing of DNA extracted from glycerol stocks carried out at GENEWIZ (South Plainfield, New Jersey, U.S.A.). Sequencing was performed in both directions using the company’s proprietary universal 16S rRNA primers, yielding assembled sequences ∼1100 nt in usable length. Species names were assigned using the Ribosomal Database Project’s Seqmatch module^28^ based on the type strain with the highest seqmatch score relative to the query strain. Three strains (*B. toyonensis* 1, 2, and 3) had identical 16S rRNA sequences, and were therefore differentiated using a 404-bp fragment of the *pyrE* gene amplified using the primers 5′-TCGCATCGCATTTATTAGAA-3′ and 5′-CCTGCTTCAAGCTCGTATG-3′ following protocols described in ref^29^. A list of the strains used, their GenBank accession numbers, competitive scores, and inferred growth parameters is given in Supplementary Table 1. For phylogenetic analysis, sequences were aligned using MUSCLE^30^ and a tree was constructed using PhyML 3.0^31,32^.

### Estimation of single-species growth parameters

The carrying capacity of each individual strain was estimated to be its optical density at 600 nm (OD_600_) in 0.2X nutrient broth after five repeated growth-dilution cycles, starting from an initial OD_600_ of 3×10^-3^. Growth curves at OD_600_ were measured in flat-bottomed 96-well microtiter plates (BD Biosciences) with lids sealed with Parafilm in a Tecan Infinite M200 Pro plate reader over 48 hr at 25⁎ C with maximum shaking. An approximation of the exponential growth rate of each individual strain was extracted from the growth curves using the time each strain took to reach a threshold optical density. The time-to-threshold method was chosen over other estimates of growth rate due to wide variations in growth patterns across the strains, which led to difficulties in fitting parameters to other population growth models.

### Competition experiments

Prior to competition experiments, cells were streaked out on nutrient agar plates, grown for 48 hr at room temperature, and then stored at 4⁎ C for up to two weeks. Single colonies were picked from these plates and grown for 24 hr at room temperature in 0.2X nutrient broth.

The competitions were initiated by diluting each individual strain in 0.2X nutrient broth to an OD_600_ of 3×10^-3^. The diluted cultures were then mixed by volume to the desired starting ratios of 0.05/0.95 and 0.95/0.05 (Strain A/Strain B). The competitions were performed in 200 ⁎L volumes in flat-bottomed 96-well microtiter plates sealed with Titer Tops® polyethylene sealing films (Diversified Biotech). For each growth-dilution cycle, the cultures were incubated at 25⁎ C and shaken at 900 rpm for 24 hr. At the end of each cycle, the cultures were thoroughly mixed and then diluted by a factor of 100 into fresh medium. OD_600_ was measured at the end of each cycle, and final species fractions were estimated after five (or, in the case of initially low plating density, seven) cycles.

To measure the final species fractions, the co-cultures were diluted by a factor of 10^4^-10^6^ (depending on OD_600_) in PBS. Seventy-five ⁎L of the diluent was plated onto 10 cm Petri dishes containing 25 mL of nutrient agar and incubated at room temperature for 48 hr. All but a small fraction of the strain pairs have distinct colony morphologies, so species fractions were estimated by counting colonies of each type (median: 51 colonies per plate). Next-generation sequencing of a subset of the co-cultures affirmed the overall accuracy of the plating technique (Supplementary Fig. 6).

### Determining the outcome of competition

The result of competition was classified as one of three outcomes: exclusion of a single strain, coexistence of both strains, or bistability. A strain was said to exclude its competitor if it was the sole strain observed from both starting frequencies after 5 cycles, or if it excluded its competitor when starting from an initial frequency of 0.95 and achieved a frequency of 0.85 or greater when starting from an initial frequency of 0.05. Pairs were considered bistable if the strain that started out at a frequency of 0.95 excluded the competitor. All other outcomes were classified as coexistence.

### Calculating competitive score and network hierarchy score

The competitive score *s*_*i*_ of each strain *i* was defined as its mean fraction *f*_*ij*_ after co-culture with each of the *n* – 1 competitor strains:

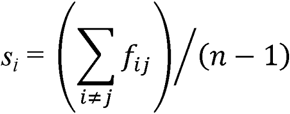

The hierarchy score (*h*) for an *n*-member network is calculated as:

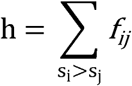

The network hierarchy score for the observed set of competitive outcomes was then compared against the distribution of scores for 10,000 simulated networks in which each pair was randomly assigned an outcome of exclusion, coexistence, or bistability with probability proportional to the incidence of each outcome in the empirical dataset. The resulting distribution of hierarchy scores was approximated using the normal distribution to determine *p-*values.

### Identifying network motifs

The frequencies of distinct topologies among the 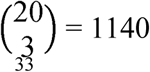 three-strain networks were enumerated using the FANMOD software package^33^. Random networks were simulated by assigning the outcome of exclusion to each pair of strains within the simulated network with the probability 0.818, which is equal to the fraction of pairs in the empirical dataset that exhibited exclusion. The occurrences of rock-paper-scissors and feedforward loop motifs were enumerated for 1000 simulated networks and approximated by a normal distribution to determine two-sided *p*-values.

### Data and code availability

The data that support the findings of this study are available from the corresponding authors upon reasonable request. An implementation of the routine for estimating the distribution of hierarchy scores and motifs in randomized networks is also available upon reasonable request.

## Acknowledgments

The authors thank S. Higgins, M. Polz, O. X. Cordero, and members of the Gore Laboratory for critical input and comments on the manuscript. This work was supported by the Defense Advanced Research Projects Agency’s Biological Robustness in Complex Settings (BRICS) program, a National Institutes of Health New Innovator Award (NIH DP2), a National Science Foundation Graduate Research Fellowship, a National Science Foundation CAREER award, a Sloan Research Fellowship, the Pew Scholars Program, and the Allen Distinguished Investigator Program.

## Author Contributions

L. M. H., J. F., and J. G. designed the study. L. M. H. performed the experiments. L. M. H., J. F., and H. S. performed the analyses. L. M. H., J. F., H. S., and J. G. wrote the manuscript.

